# Development of an Escape-resistant SARS CoV-2 Neutralizing Synthetic Nanobody

**DOI:** 10.1101/2022.06.07.495065

**Authors:** Dmitri Dormeshkin, Michail Shapira, Simon Dubovik, Anton Kavaleuski, Mikalai Katsin, Alexandr Migas, Alexander Meleshko, Sergei Semyonov

**Affiliations:** Institute of bioorganic Chemistry of the National Academy of Sciences of Belarus, Minsk, Belarus; Belarusian State University, Minsk, Belarus; Institute of Science and Technology Austria, Klosterneuburg, Austria; Imunovakcina, UAB, Vilnius, Lithuania; Imuunofusion, LLC, Minsk, Belarus; Republican Research and Practical Center for Epidemiology & Microbiology, Minsk, Belarus

**Keywords:** COVID-19, Nanobodies, SARS-CoV-2, Synthetic library, Neutralization, RBD, VHH

## Abstract

An emerging COVID-19 pandemic resulted in a global crisis, but also accelerated vaccine development and antibody discovery. In this work, we identified a number of nanomolar-range affinity VHH binders to SARS-CoV-2 variants of concern (VoC) receptor binding domains (RBD), by screening synthetic humanized antibody library with more than 10^11^ diversity. In order to explore the most robust and fast method for affinity improvement, we performed affinity maturation by CDR1 and CDR2 shuffling and avidity engineering by multivalent trimeric VHH fusion protein construction. As a result, H7-Fc and G12×3-Fc binders were developed with the affinities in nM and pM range respectively. Importantly, their affinities are weakly influenced by SARS-CoV-2 VoC mutations. The plaque reduction neutralization test (PRNT) resulted in IC50 = 100 ng\ml and 9.6 ng\ml for H7-Fc and G12×3-Fc antibodies respectively for emerging Omicron variant. Therefore, these VHH could expand the present landscape of SARS-CoV-2 neutralization binders with the therapeutic potential for present and future SARS-CoV-2 variants.

## Introduction

The current emerging severe acute respiratory syndrome coronavirus 2 (SARS-CoV-2) causes a global pandemic and the coronavirus disease (COVID-19)-related deaths had exceeded 5.6 million in February 2022 (1). Despite the unprecedented success in SARS-CoV-2 vaccine development this number is still growing, raising the question of multiple antiviral therapeutics development.

The SARS-CoV-2 virus entry is initiated on the host cell surface by the attachment of spike (S) protein receptor-binding domain (RBD) to angiotensin converting enzyme 2 (ACE2). It became obvious that antibodies could prevent virus fusion with the cell membrane by blocking RBD-ACE2 interaction or fixing the RBD in “down” conformation (2),(3).

Therapeutic neutralizing antibodies constitute a key short-to-medium term approach to tackle COVID-19. The last outbreak of the Omicron (B.1.1529) variant abrogated the majority of the FDA-authorized antibody treatments (4). Despite the lower mortality rate from the Omicron variant, its antibody treatment is still in high demand, especially for immunosuppressed patients, as well as high risk group patients being not able to be fully vaccinated (5). As the antibody discovery and clinical approval is a time-consuming process even under the special conditions for COVID-19 treatment authorization, it is necessary to have a pipeline of VoC neutralizing binders identification and isolation. Synthetic libraries and *in vitro* display selection processes proved themselves during the pandemic yielding a variety of SARS-COV-2 antibodies of different origins and shapes in quick terms (6),(7),(3).

In this study, we report a pipeline for the generation of VHH-based neutralizing binders from diverse humanized synthetic libraries. We have identified an antibody G12, that could bind Wuhan-Hu-1 (WT) and Delta (B.1.617.2) variants of SARS-CoV-2 with the two-digits nanomolar affinity constant. Utilizing affinity and avidity engineering we explored the possibility of therapeutic potential enhancement of this clone. We succeeded in generation of affinity maturated H7-Fc with 2-9 nM affinities range to VoCs including Omicron and triple G12×3-Fc with the pM affinity to the Delta variant and 3 nM affinity against Omicron. We have demonstrated that isolated antibodies possess strong binding and neutralization characteristics for highly transmissible Delta and Omicron variants, not inferior and even superior to FDA approved (EUA) antibodies for COVID-19 treatment.

This selection strategy could be efficiently expanded towards new variants of SARS-CoV-2 in order to expand the currently existing landscape of VHH neutralizing binders with the therapeutic potential. The data suggest that H7-Fc and G12×3-Fc antibodies could be a promising therapeutic agent for COVID-19 treatment, which may retain efficacy against continuously developing variants.

## Results

### Identification of VHH against SARS-CoV-2

The synthetic humanized nanobodies library VHH4.0_DD was used as a source of RBD binders. This library consists of 4 sub-libraries with a diversity of 2-3×10^10^ clones each. Each sub-library has a tailored diversity of CDRs of different lengths mimicking the natural *Camelid* antibodies diversity, but exceeding it in its size. The VHH4.0_DD library proved its suitability previously yielded a number of low-nanomolar binders to different targets, including difficult poly-glycosylated (unpublished data).

To enrich binders that block the interaction of RBD with the ACE2 receptor we have utilized a negative selection approach (Figure 1).

**Figure 1.**
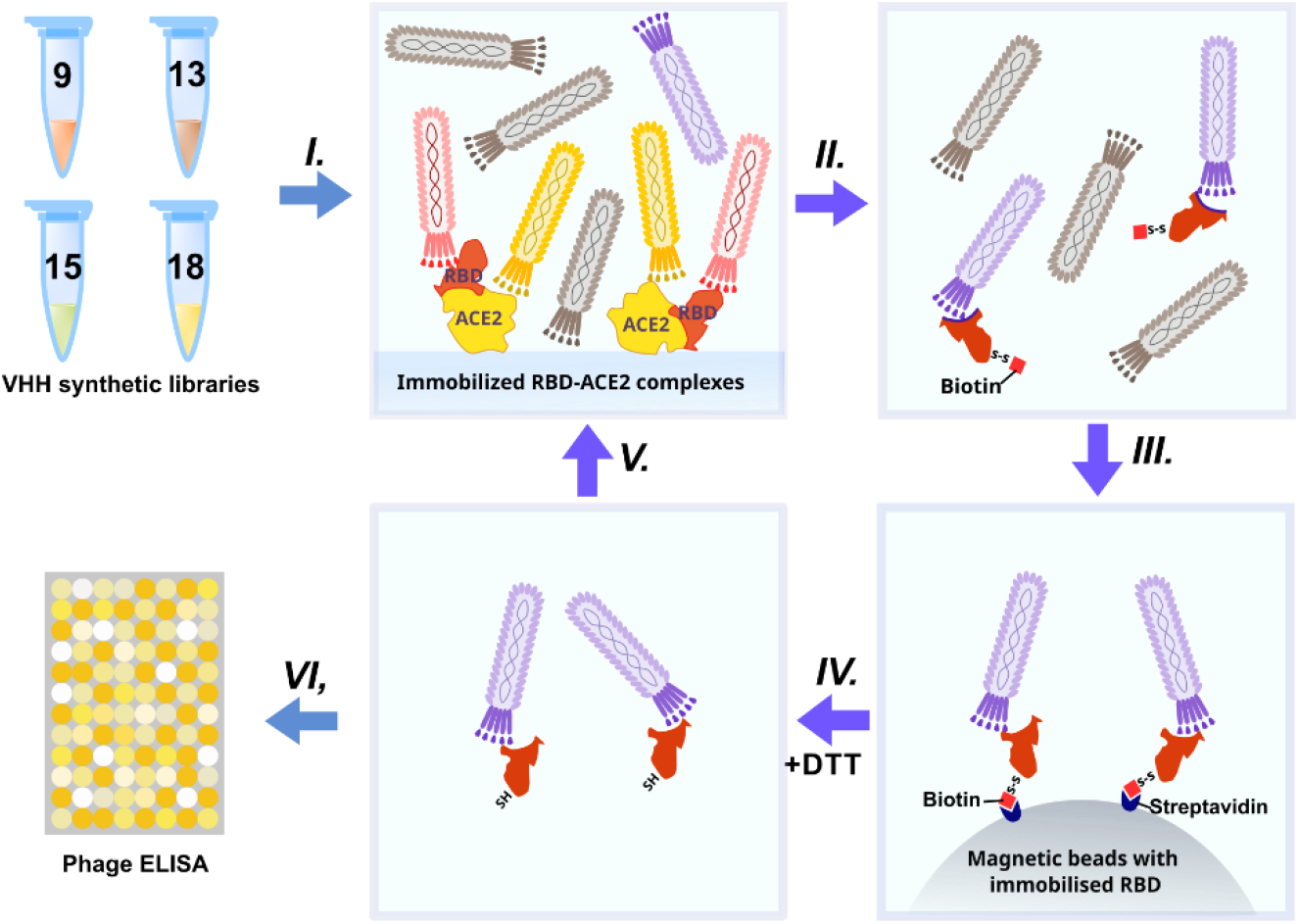
Schematic representation of RBD-binders phage display selection.

Briefly – phage display libraries were incubated with the immobilized RBD-ACE2 complex (*I)*. Binders that bind the RBD-ACE2 interface mostly remained in the solution while sticky and unspecific binders were depleted. Then the supernatant was transferred to an Eppendorf tube with the biotinylated RBD protein (*II*) (WT for the first round, Beta (B.1.351) for the second, Delta for the third, and Omicron for the fourth) for 2 h incubation and pulled down with the streptavidin magnetic beads afterward *(III)*. As RBD proteins were chemically biotinylated with the -S-S-containing agent, mostly RBD-specific phages were eluted with DTT treatment from magnetic beads after pull down and washing *(IV)*. Off-rate selection during the last round ensured the selection of the tightest binders with the slowest dissociation rate (*k*_*off*_) and the highest affinity (8). The concentration of the RBD was sequentially reduced from 200 nM in the first round to 10 nM in the last round *(V)*.

After four rounds of selection 32 individual clones were analyzed by monoclonal ELISA (figure 2A). The binding of each clone to the RBD was compared with the binding to a BSA, resulting in a signal-to-noise ratio >10 for 20\32 clones.

**Figure 2.**
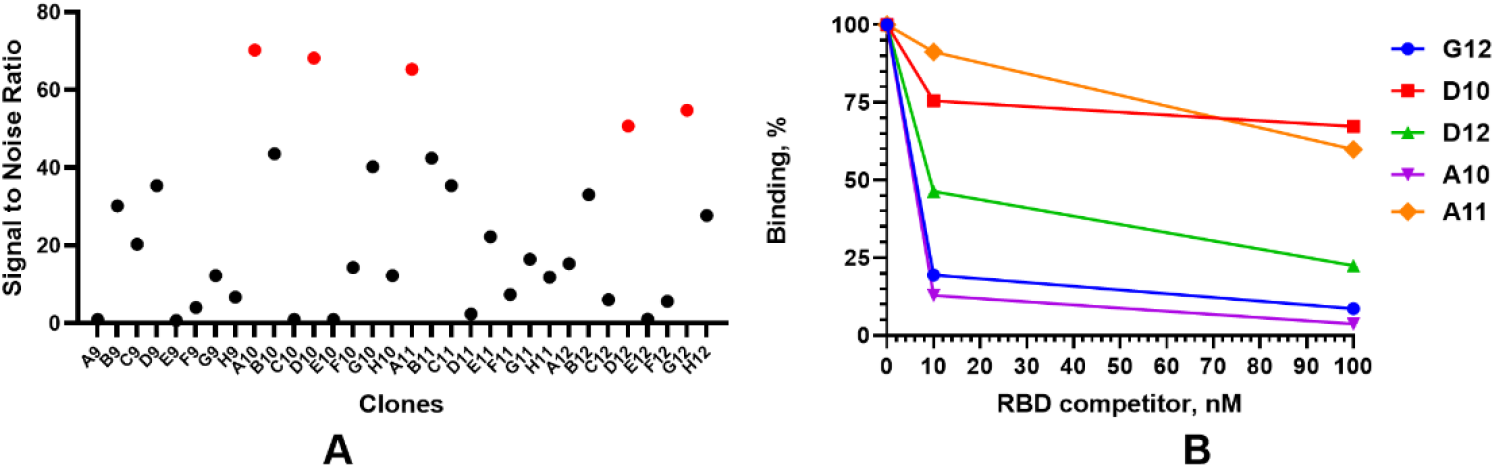
Monoclonal ELISA of selected VHH clone (A) and competitive phage ELISA (B)

A competitive phage ELISA allowed us to range the top 5 binders by their *k*_off_value and exclude binders that poorly bind soluble antigen in a liquid phase (Fig. 2B). Antibodies with the fastest dissociation rate were recaptured by the immobilized antigen resulting in a high binding signal after anti-M13-HRP detection. The steepest signal decreasing with the increase of the antigen concentration (from 0 to 100 nM) corresponded to the binders with the slowest dissociation. Antibodies with the slowest dissociation rate are considered better binders.

The most promising G12 and A10 binders were reformatted in the VHH-Fc fusion format for affinity and specificity determination against WT and Delta strains by means of biolayer interferometry using Octet R2 (Figure 3).

**Figure 3.**
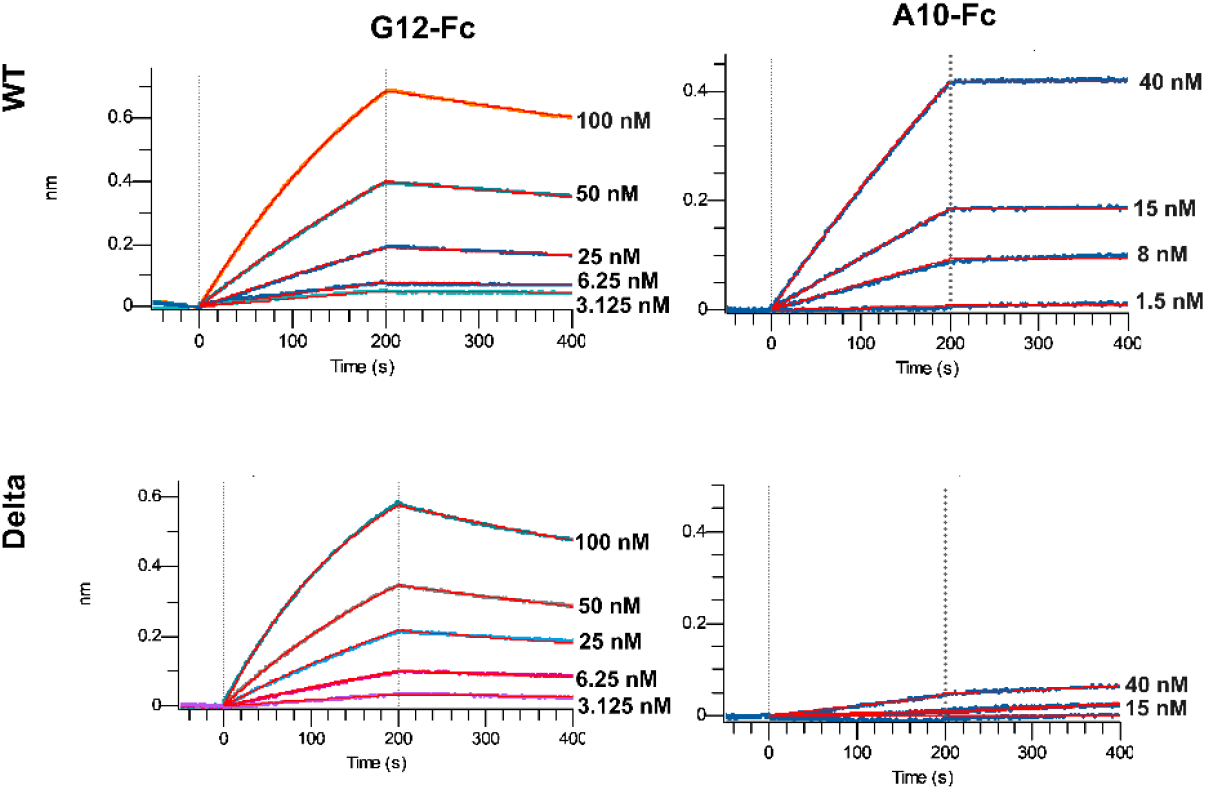
Multipoint Biolayer Interferometry (BLI) of VHH-Fc binders to WT and Delta variants RBD.

The A10-Fc antibody possessed a very high binding affinity of 25 pM, but showed no significant binding to the Delta. Antibody G12-Fc had a lower affinity of 20 nM, but its binding was not affected by Delta variant’s mutations. Two digits nM affinity of the G12-Fc allows it to be considered as a high-affinity binder, but it could not be sufficient for effective neutralization of SARS-CoV-2 infection with the IC50 in a sub-nanomolar range.

In order to enhance the binding affinity of G12 we utilized in parallel two strategies – avidity enhancement by trivalent tandem repeats VHHx3-Fc protein construction and affinity maturation of G12 antibody by CDR1 and CDR2 shuffling.

### Affinity engineering of G12 binder

Affinity maturation is a well-established approach for antibody binding properties enhancement (9). In the absence of VHH-RBD complex structure, there are two options left to choose between – error-prone PCR and CDR shuffling. CDR3 contributes the most in VHH interactions with antigens, so we decided to re-randomize CDR1 and CDR2 loops to not compromise affinity and specificity that we already had.

In our library VHH4_DD unique restriction site was incorporated after the CDR2 region in order to provide the possibility of FR1-CDR2 region randomization by simple restriction\ligation steps. *E*.*coli* colonies harboring the VHH4_DD library were scrapped from the 2 large 200 mm plates and were used for the phagemids pool preparation. FR1-CDR2 region from this pool was subcloned into G12 VHH for the secondary library formation. The resulting diversity of the secondary library was measured by the transformants counting after the electroporation and was 10^8^ clones.

Three rounds of the selection were performed with the biotinylated RBD concentration decreasing from 50 nM to 1 nM. After the third round monoclonal ELISA revealed that 15 out of 16 clones bind RBD with >10 signal-to-noise ratio (Figure S1). Sequence alignment showed a single sequence H7 prevalence which was presented in 12 from the 15 sequences. For the affinity measurements H7 clone was reformatted in the VHH-Fc fusion format and purified from the transiently transfected HEK293 FreeStyle supernatant (Figure S2A).

The binding analysis showed a 5.57-fold affinity improvement for H7 with the 4.4 nM *K*_*D*_ value for the interaction with WT SARS-CoV-2 variant. We also performed extended binding studies with the emerging VoC RBDs – Delta, Delta plus, Beta and Omicron (Figure 4A).

**Figure 4.**
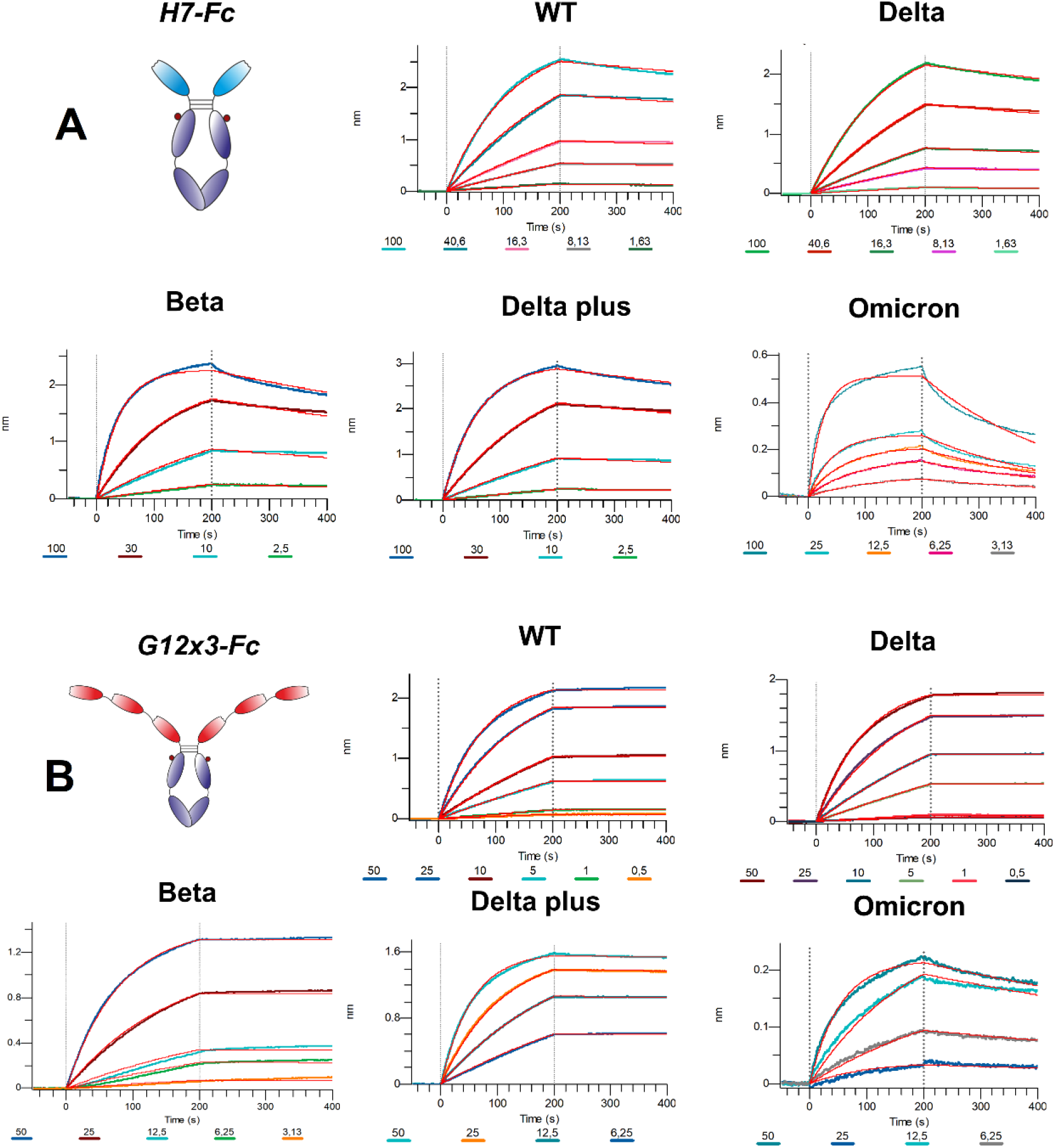
Multipoint BLI of H7-Fc (A) and G12×3-Fc (B) binders to SARS-CoV-2 VoC RBDs.

It was revealed that VoC mutations slightly affect the affinity of H7 to different RBDs and its *K*_*D*_value remains in the single-digit nanomolar range (Table 1).

**Table 1.** Antibodies binding activity against RBD variants; related to Figure 4.

|  | G12x3-Fc |  |  |  | H7-Fc |  |  |  |
| --- | --- | --- | --- | --- | --- | --- | --- | --- |
| | $K_D$ (M) | $k_{on}$ (1/Ms) | $k_{off}$ (1/s) | $R^2$ | $K_D$ (M) | $k_{on}$ (1/Ms) | $k_{off}$ (1/s) | $R^2$ |
| **WT** | $<1 \times 10^{-12}$ | $2.624 \times 10^5$ | $<1 \times 10^{-7}$ | 0.9996 | $4.485 \times 10^{-9}$ | $8.937 \times 10^4$ | $4.008 \times 10^{-4}$ | 0.9991 |
| **Delta** | $<1 \times 10^{-12}$ | $2.413 \times 10^5$ | $<1 \times 10^{-7}$ | 0.9996 | $7.722 \times 10^{-9}$ | $7.213 \times 10^4$ | $5.570 \times 10^{-4}$ | 0.9995 |
| **Delta+** | $1.562 \times 10^{-10}$ | $3.688 \times 10^5$ | $5.760 \times 10^{-5}$ | 0.9993 | $3.414 \times 10^{-9}$ | $1.634 \times 10^5$ | $5.579 \times 10^{-4}$ | 0.9995 |
| **Beta** | $<1 \times 10^{-12}$ | $2.538 \times 10^5$ | $1.226 \times 10^{-7}$ | 0.9983 | $3.722 \times 10^{-9}$ | $2.535 \times 10^5$ | $9.435 \times 10^{-4}$ | 0.9976 |
| **Omicron** | $2.968 \times 10^{-9}$ | $3.571 \times 10^5$ | $1.060 \times 10^{-3}$ | 0.9944 | $9.493 \times 10^{-9}$ | $4.342 \times 10^5$ | $4.122 \times 10^{-3}$ | 0.9883 |

Considering the broad binding spectrum against SARS-CoV-2 variants it was decided to improve its binding properties and a neutralization potency by another approach – avidity enhancement.

### Avidity engineering of G12 binder

One of the remarkable features of the VHH binding domains is that they could be assembled into multivalent structures with the non-linear increase of the binding properties (10). In addition to the expected avidity effect, it was assumed that linked VHH domains could bridge the distance between different RBDs on the same S-protein, but not between different S-proteins because of the inter-S-protein distance on the viral surface (11). It allows us to assume a superiority of the tandem VHH variant in a live virus neutralization test.

A multivalent trimeric clone G12×3-Fc was constructed by fusion of three G12 sequences through flexible linkers L1 (GGGGSGGGGSGGGGS) and L2 (GGGGSGGGGSSGGGS) with the Fc domain of IgG1 protein. The resulting protein was transiently expressed in HEK293 FreeStyle and purified to the homogenous state (Figure S2B). Expression level decreased insignificantly from 15 mg\l to 12 mg\l.

Affinity measurements were performed by BLI on Octet R2 using WT and VoC RBDs (Figure 5B). Kinetic data comparison with the affinity maturated H7-Fc protein revealed a 2-fold increase of *k*_*on*_ values and more than 100-fold increase of *k*_*off*_ values for WT, Beta, Delta and Delta plus variants (Table 1). Surprisingly, G12×3-Fc binding to Omicron increased unproportionally only to 2.968 nM.

**Figure 5.**
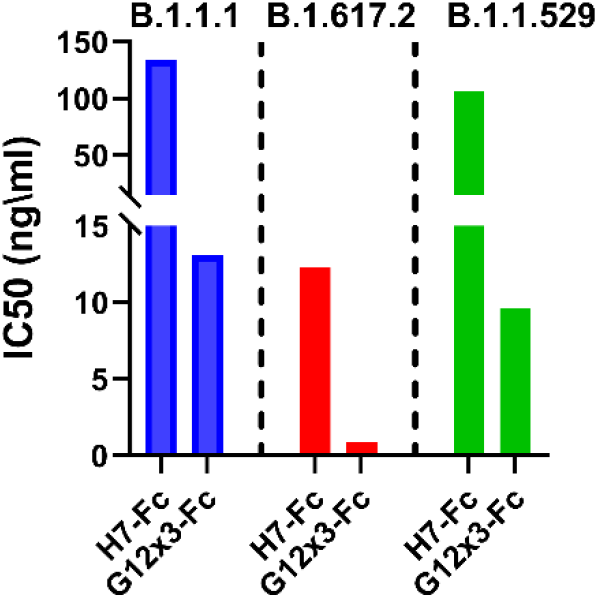
Half-maximal inhibitory concentrations (IC50, ng\ml) values chart for H7-Fc and G12×3-Fc against SARS-CoV-2 variants.

We have also engineered G12×2-Fc tandem repeat fusion protein with the affinity to Delta and Beta variants of 0.6 nM and 1.6 nM respectively (Figure S3). It was considerably less than the G12×3-Fc picomolar affinity, therefore this clone was not considered further.

Nevertheless, the most important criterion for the SARS-CoV-2 antibody potency is its neutralization, quantified by the inhibitory concentration (IC) values (e.g. IC50).

### Neutralization assay

The state-of-the-art method of determining the neutralization potential of antibodies is the plaque reduction neutralization test (PRNT). In this method, Vero E6 cells are infected with the different variants of SARS-CoV-2 in the presence of serially diluted antibodies (12).We used PRNT to determine if the H7-Fc and G12×3-Fc proteins were able to neutralize B.1.1.1, Delta and Omicron SARS-CoV-2 virus variants. The neutralization IC50 potencies of these antibodies are shown in Figure 5.

As the results of virus neutralization assays could vary from test to test in various laboratories we included ACE2-Fc protein with the well-known IC50 value for different variants (Table S1).

H7-Fc antibody neutralized SARS-CoV-2 strains with an IC50 in a 12.3-133.8 ng\ml range. Encouragingly, the Omicron variant could not escape the neutralization. G12×3-Fc binder showed a 10-fold decrease of IC50 values with 13.1 ng\ml, 0.9 ng\ml, and 9.6 ng\ml for B.1.1.1, Delta and Omicron respectively.

## Discussion

Monoclonal antibodies due to their immediate virus neutralization properties represent a promising strategy for SARS-COV-2 treatment (13). Despite the lowering of a number of patients with severe cases in 2022, people who have immunosuppression, including transplant recipients and people with cancer, advanced or untreated HIV and autoimmune disorders as well as patients with high risk features (older age>65, obesity or being overweight, pregnancy, chronic kidney disease, diabetes mellitus, cardiovascular disease, hypertension, chronic lung disease) still need the antibody treatment and pre- or post-exposure prophylaxis (5).

At the time of manuscript preparation, the Omicron variant had become the dominant strain due to the highest transmissibility and the possibility to evade humoral immune response induced by all the major vaccines (14). Most of FDA approved therapeutic monoclonal antibodies including REGN10933, REG10987, LY-CoV555, LY-CoV016, AZD8895, and AZD1061 were found to be ineffective against the Omicron BA.1/BA.2 variants, while AZD7442 (Tixagevimab–Cilgavimab) and Sotrovimab were partially effective (15), (16), (17). Moreover, it was suggested that Sotrovimab (and others that form the few monoclonal antibodies, that are potentially effective against Omicron) could drive SARS-CoV-2 escape in immunocompromised patients (18). A wide variety of available antibody treatment options, including cocktails, or monoclonal antibodies rotation could alleviate this problem.

In this work, we explored the possibility of the isolating of promising VHH binders with the desired properties from the synthetic VHH libraries. Such an approach has been proven to be successful by Seeger’s Sybodies libraries (19) and Kossiakoff’s synthetic high-performance Fab library (7), but with no bias towards variants of concern.

Our objective was to generate a number of SARS-CoV-2 neutralization binders that could tolerate most of the VoC mutations by selection strategy modification. It was previously reported that the depletion of anti-RBD antibodies in convalescent patient sera results in the loss of >90% neutralizing activity of these sera against SARS-CoV-2 (20). It suggests that RBD binding by IgGs is the main and the most efficient mechanism of SARS-CoV-2 neutralization. It also has been discovered, that RBD is the most potent immunogen protein for *Llama spp*. immunization for VHH generation in comparison with the S-protein or S1-subunit (21).

The small size and single-domain nature of VHH allow them to bind epitopes that are not available for conventional antibodies, in particular concave epitopes such as grooves and clefts (22). Synthetic origin of the library also contributes to an increase in the possible diversity of epitopes compared to human antibodies.

During the first round we used the whole spike protein trimer as an immunogen to guide the selection towards the selection of antibodies against epitopes located within the RBD in the correct conformation of the intact spike protein trimer. For every subsequent round a different RBD variant – Wuhan-Hu-1 (WT), B.1.359 (Beta), B.1.617.2 (Delta) and B.1.1.529 (Omicron) as an antigen was used. Besides, negative selection using RBD-ACE2 complexes was performed in order to deplete binders that do not interfere with RBD-ACE2 binding. This is one of the key advantages of synthetic libraries – a precise control under the selection pressure, including negative selection and antigens rotation.

A panel of RBD specific VHH was selected with a picomolar and nanomolar binders among them. It was revealed that all binders have different sequences with CDR3 lengths of 9 and 13 amino acids. Despite the fact that the pool of libraries contains sub-libraries with CDR3 lengths of 9, 13, 15, and 18 amino acids, the longest CDR3 sequences were not enriched during the selection. Presumably shorter CDR3 is a more favorable paratope shape which is more suitable for the binding on the RBD-ACE2 interface.

The potent antibody G12-Fc was isolated with the 20 nM affinities against WT and Delta variants. Two strategies not exceeding 3 weeks in length were tested to boost its potency.

In the absence of prior structural investigation, it was decided to perform affinity maturation by a blind CDR1 and CDR2 randomization. During the library construction step, we have included restriction sites flanking CDR1-CDR2 region in order to clone these CDRs diversity in selected candidates’ sequences. Isolated from the secondary affinity maturated library clone H7 demonstrated the 5.57-fold affinity improvement with the 4.4 nM *K*_*D*_ value for the interaction with WT SARS-CoV-2 variant. Its affinities against Delta, Delta plus, Beta, and Omicron variants were 7.2 nM, 3.4 nM, 3.7 nM, and 9.5 nM respectively.

It was suggested, that the H7 clone belongs to the most promising class 3 or class 4 binders that prevent the Omicron escape (23). Nevertheless, affinity improvement of H7 was quite low in comparison with successful cases of affinity increasing by more than two orders of magnitude (24). A modest affinity enhancement could be explained by the non-significant role of CDR1 and CDR2 loops of G12 antibody in the interaction with RBD. This is not an uncommon situation for single-domain antibodies – to interact predominantly or even solely by the CDR3 hypervariable loop (25).

A multivalent trimeric clone G12×3-Fc was constructed with superior to G12-Fc and H7-Fc antibodies binding properties. *K*_*D*_ values in a picomolar range for WT, Beta, Delta and Delta plus variants put it among the most tightest RBD binders (26).

We investigated the neutralization activity of H7-Fc and G12×3-Fc antibodies using the plaque reduction neutralization test against live SARS-CoV-2 VoCs. It was elucidated that both of the antibodies efficiently block the virus entry into cells even regardless of the Omicron escape mutations. Despite the significant difference in binding constants, neutralization potency of G12×3-Fc and H7-Fc against live SARS-CoV-2 differed by no more than a factor of 10 for different VoCs.

Nevertheless, G12×3-Fc neutralization potency is not inferior and even superior to most of the SARS-CoV-2 binders approved to date, which proves the power of large synthetic antibody libraries and sophisticated selection techniques for cost-time effective antibodies with the therapeutic potential isolation (27).

The antibodies described in this report could expand the current landscape of therapeutic COVID-19 antibodies as well as being used for elucidation and clarification of the structural mechanisms of virus neutralization. As the synthetic antibodies selection and design strategy described above significantly differs from the natural immune response it could lead to more diverse paratope space and results in more promising binders not only for SARS-CoV-2 but also for other antigens with the high mutation rate.

## Material and Methods

### Reagents and solutions

Sulfo-NHS-SS-Biotin, streptavidin-PolyHRP, Dynabeads T1, Tween-20, PEG8000, FreeStyle™ Expression Medium, Nunc Maxisorp high-binding plates were purchased from Thermo Fisher Scientific. TB, LB, 2xYT medium, PBS were bought from Melford Laboratories Ltd. M13KO7 phage helper, *ER2738* strain and all enzymes were purchased from New England Biolabs Inc. Other chemicals were purchased from Sigma and used without further purification and additional preparation. All solutions were prepared using deionized MilliQ quality water.

### SARS-CoV-2 virus variants

Three isolates of SARS-CoV-2 were used: 2245 (B.1.1.1), 2107 (B.1.617.2.122, Delta derivative), 1984 (B.1.1.529.1, Omicron variant). The isolates were obtained in 2021 and 2022 at the Biosafety laboratory of the Republican Scientific and Practical Center for Epidemiology and Microbiology (Republic of Belarus) from nasopharyngeal swabs of patients with symptoms of COVID-19 and typed using sequencing.

### Production of Recombinant SARS-CoV-2 RBD variants and hACE2 protein

For the initial VHH selection experiments and binding affinity measurements a codon optimized RBD (residues 319 to 529) of spike protein from SARS-CoV-2/human/China/Wuhan-Hu-1/2019 (GenBank: QHD43416.1) and Omicron (IPBCAMS-OM01/2021) were generated by gene synthesis (Synbio-tech). Selected mutations for the generation of Beta B.1.351, Delta B.1.617.2 and Delta plus B.1.617.2/AY.1 variants were introduced into Wuhan-Hu-1 RBD by site-directed mutagenesis with the primers listed in Table S2. All the RBD variants were cloned into mammalian expression vector pcDNA3.2 (Thermo Fisher) with the additional N-terminal HA leader sequence (MNTQILVFALIAIIPTNADKIGSGA) and C-terminal 10x His tag. ACE2 gene was amplified from cDNA with the primers hACE2-f\hACE2-r and cloned into a custom modified pFUSE-hIg1d expression vector (Invitrogen).

Endotoxin-free purified plasmids were transiently transfected in FreeStyle™ 293-F cells. FreeStyle™ 293-F cells were passaged at 0.5 × 10^6^ cells/mL in 40 mL of fresh FreeStyle™ Expression Medium in 250 ml polycarbonate shake flask (Corning). The following day, 40 μg of DNA and 80 μL of polyethyleneimine (10 kDa, linear), separately diluted in 0.4 mL of Opti MEM (Gibco), were vigorously mixed. The mixture was incubated for 25 minutes at RT, and then added to cells. Transfected cells were incubated at 37 °C, 8% CO2 for 5 days on an orbital shaker platform (125 rpm). Transfected supernatants were collected 5 days after expression, clarified with centrifugation (800 g, 5 min), filtered with a 0.2 μm PVDF filter and purified over HisPur and Protein A column for RBD and hACE2-Fc proteins respectively using ÄKTA Purifier 10 FPLC System (Cytiva Life Sciences). Both recombinant SARS-CoV-2 RBD and hACE2-FC were further purified to homogeneity using a Superdex 75 Increase 10/300 column (Cytiva Life Sciences).

The RBDs were biotinylated by amine coupling with sulfo-NHS-SS-Biotin according to the manufacturer’s manual (Thermo Fisher). Briefly, fresh sulfo-NHS-SS-Biotin (10 mg/mL in PBS; pH 7.4) was rapidly added to RBD in PBS at a molar ratio of 20:1. After 1 h of reaction at room temperature (RT) with gentle shaking, free biotin was removed through extensive dialysis at 4°C. Then biotinylated RBDs were stored in aliquots (at -80°C until use).

The full-size SARS-CoV-2 protein was purchased from the commercial vendor Invitrogen (#RP-87680).

### Identification of SARS-CoV-2 binders

A proprietary humanized synthetic VHH library with a diversity exceeding 10^11^ unique antibody clones was used as a source of RBD-specific binders. Two plates of high-binding Nunc microplate wells were coated with hACE2-FC at 2 μg/ml in PBS overnight at 4°C (100 μl per well). The next day plate was blocked with 1% BSA in PBST for 1 h and 3 ug/ml RBD was added for 1 hour. At the same time 20 μl of Dynabeads T1 streptavidin magnetic beads were blocked with 0.5% BSA for 1 h and then washed 3 times with 0.1% PBST (PBS supplemented with 0.1% Tween-20). VHH libraries were added to RBD-ACE2 complexes for 30 min and then were transferred to a 1.5 ml low protein-binding tube with 100 nM biotinylated RBD in 0.5% BSA. After 60 min of incubation, the phages/RBD mixture was transferred to streptavidin magnetic beads for 10 min. Then the tube was placed in a magnetic particles separator and allowed to stand for 30 s to remove the supernatant containing the unbound phages. The particles were washed 10 times by cycles of magnetic rack separation and resuspension in PBST. Finally, the phages were released by 30 min incubation with 100 mM DTT for -S-S-bond disruption and specific elution of RBD-bounded phages. Eluted phages were used for titration and amplification in *E. coli ER2738* for additional rounds of selection. In total four rounds of selection were made. Selective pressure was maintained by RBD concentration decreasing (from 100 nM to 10 nM) and washing stringency increasing (from 10 times in the 1st round to 20 times in the 4th). In the last round, off-rate selection with the presence of 5 μM of non-biotinylated RBD was performed within 5 min. After the last round, individual clones from the titration plate were used for phage supernatants production by overnight cultivation in 200 μl of 2xYT medium, supplemented with 50 μg/ml kanamycin and 100 μg/ml carbenicillin, in 96-well cell culture plate at 30°C with continuous shacking. Cells were pelleted at 3200 g for 30 min. Twenty μl of 10x PBS with 0.5% NaN3 was added to phage supernatants before storage at 4°C for phage ELISA.

### Phage monoclonal and competitive ELISA

For phage monoclonal ELISA 32 wells of a 96-well microtiter plate were coated with 2 μg/ml RBD (positive wells) in 100 μl PBS and 32 wells with 5 μg/ml BSA (negative wells) overnight at 4 °C. Afterward, the plate was blocked with 5% skimmed milk in PBS for 2 hours at RT. Five μl of individual phage supernatants were mixed with 95 μl of 2.5% (w/v) skimmed milk. The mixtures were added to the wells and incubated at RT for 30 min. After incubation, the wells were washed six times with 0.1% PBST and 100 μL of anti-M13 phage antibody conjugated with HRP (1:5000 dilution in PBS) was added. After 1 h of incubation and six washes, positive binders were determined upon 100 μl TMB substrate addition. The absorbance at 450 nm was determined after stopping the reaction by adding 100 μL of 2 M H_2_SO_4_ per well. The top twenty binders according to their signal-to-noise ratio were subjected to sequencing and a competitive ELISA assay.

For the competitive phage ELISA, a RBD-coated plate was prepared as described before. Phages (10^9^ PFU/ml) in 100 μl of 2.5% skimmed milk were incubated with three different concentrations of RBD (0 nM, 10 nM, 100 nM) for 1 hour at RT with continuous shacking. Then mixtures were transferred to RBD-coated wells for 15 min and washed six times with 0.1% PBST. Detection of bounded phages was done as described before for monoclonal ELISA.

### VHH affinity maturation and tandem repeats formation

For the affinity maturation of the G12 clone FR1-CDR1-FR-2-CDR2 region was randomised by subcloning this region from pooled synthetic libraries phagemids into the G12 sequence. The resulted diversity was electroporated into *E*.*coli ER2738* pre-infected with M13KO7 phage for G12-based library construction as described elsewhere (28). Three rounds of biopanning were performed with different RBD concentration from 10 nM to 100 pM as described above for SARS-CoV-2 binders identification.

For multimeric tandem VHH construction G12 was amplified in three separate PCR reactions with primers pairs VHHtri-NcoI-f/VHHtri-BamHI-r, VHHtri-BamHI-f/VHHtri_XhoI-r and VHHtri-XhoI-f/VHHtri-NotI-r (Table S2). VHH fragments were digested by *NcoI\XhoI, XhoI\SalI, SalI\NotI* respectively and ligated simultaneously into *NcoI\NotI* digested into a custom modified pFUSE-hIg1 expression vector (Invitrogen).

### Production of VHHs in mammalian cells in fusion with human IgG1 Fc (VHH-Fcs)

The antibody sequences (monomeric and multimeric tandem repeats) were cloned into the pFUSE-hIg1d expression vector for transient transfection. pFUSE-hIg1d is a modified pFUSE-hIg1 (Invitrogen) in which *NotI* restriction site in the backbone was deleted by site-directed mutagenesis and incorporated before the Fc domain. The Fc-fusion proteins were produced by transient transfection of FreeStyle™ 293-F cells followed by Protein A affinity chromatography as previously described for ACE2-Fc protein. Then Fc-fusion proteins were dialyzed against PBS and concentrated with Amicon Ultra-4 centrifugal unit (MWCO 30 kDa). Their purity and integrity were verified by reducing SDS-PAGE and MALDI-TOF MS.

### Bio-layer interferometry (BLI) assay

The binding kinetics of antibodies to RBD was measured by BLI on an Octet-R2 (Sartorius). The his-tagged RBD variants were loaded onto Ni-NTA biosensors from 10 ug\ml solution in kinetic buffer (PBS, 0.02% tween-20, 0.05% BSA, filter-sterilized) for 300 s. The sensors were equilibrated (baseline) for 60 s, before incubating with VHH-Fc nanobody fusion proteins at various concentrations from 0.5 nM to 100 nM (association) for 200 s. Dissociation kinetics were measured by dipping sensor tips into wells containing kinetic buffer for 200 s. Data were reference subtracted (reference sensor and reference sample), and kinetics were calculated in Octet Analysis Studio v.12.2 using a 1:1 binding model.

### Plaque reduction neutralization test

Biosafety Level 3 laboratory setting was used for PRNT tests. For each test, serial ten-fold dilutions of samples starting at a concentration of 10 μg/ml were prepared in Dulbecco modified Eagle medium (DMEM). The dilutions, mixed to a 1:1 ratio with a virus solution containing about 30 plaque-forming units (PFU) of SARS-CoV-2 per well (previously determined by virus titration), were incubated for 1 h at 37 °C. Sixty microliters of the virus-sample mixtures were added to confluent monolayers of Vero E6 cells in 96-well plates in triplicates and incubated for 1 h at 37 °C in a 5% CO2 incubator. The inoculum was removed and 100 ml of overlay solution of DMEM, 2% fetal bovine serum (FBS), penicillin (100 U/ml), streptomycin (100 U/ml) and 1% carboxymethylcellulose were added to each well. After incubation at 37ºC in a 5% CO2 atmosphere for 76 hours, cells were fixed with a 4% formaldehyde solution, stained with crystal violet (0.05%) and plaques were counted. IC50 value was calculated by logistic regression using an online tool “Quest Graph™ IC50 Calculator.” AAT Bioquest, Inc., 21 Apr. 2022, https://www.aatbio.com/tools/ic50-calculator.

## AUTHOR CONTRIBUTIONS

Conception and design of the experiments: DD and MK. Synthetic antibody libraries assembly: DD and AMe. Phage display and affinity maturation DD. Cloning, production, purification and characterization of VHH and their Fc-fusions: MS, AK and SD. Production and purification of recombinant RBD and ACE2 proteins: AK and SD. BLI kinetic measurements DD. SARS-CoV-2 neutralization tests: AMi and SS. Data analysis and interpretation, figures preparation: DD and MS. Drafting the manuscript: DD. All authors contributed to the article and approved the submitted version.

## DISCLOSURE STATEMENT

This research is sponsored by Immunofusion, and authors MK, AMi and AMe are employees of UAB Imunovakcina and LLC Immunofusion. DD is a member of the Scientific Advisory Board of Imunovakcina.

## FUNDING

The author(s) reported there is no funding associated with the work featured in this article.

## COMPETING INTERESTS

DD and MS have a pending patent application for the RBD-targeted antibodies from this study.

## Supplementary Information

**Figure S1.**
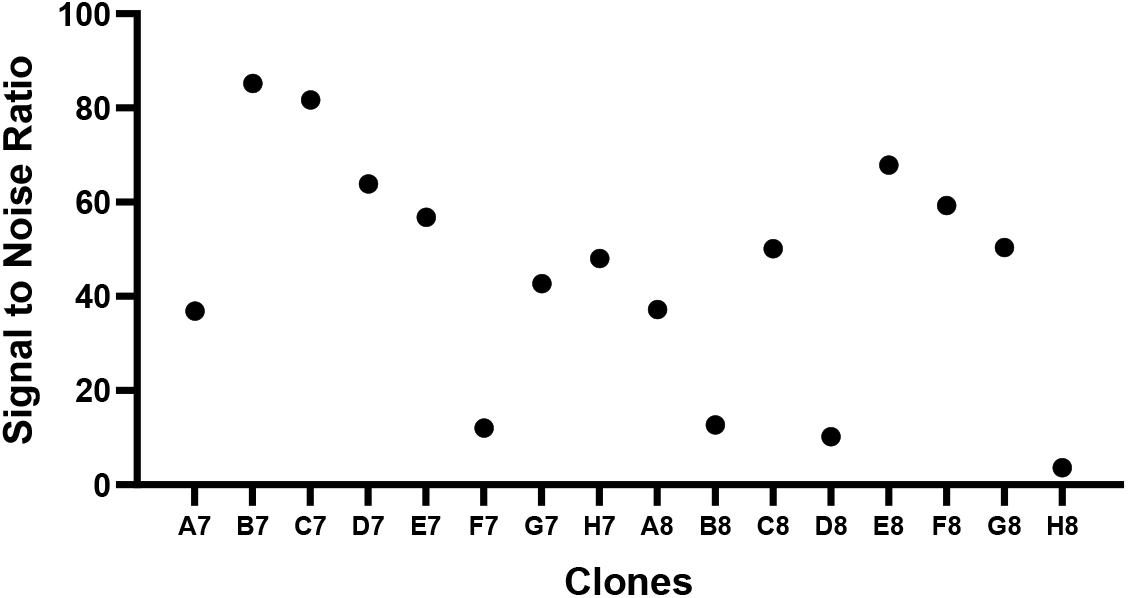
Monoclonal ELISA of selected VHH clones from affinity maturated secondary library.

**Figure S2.**
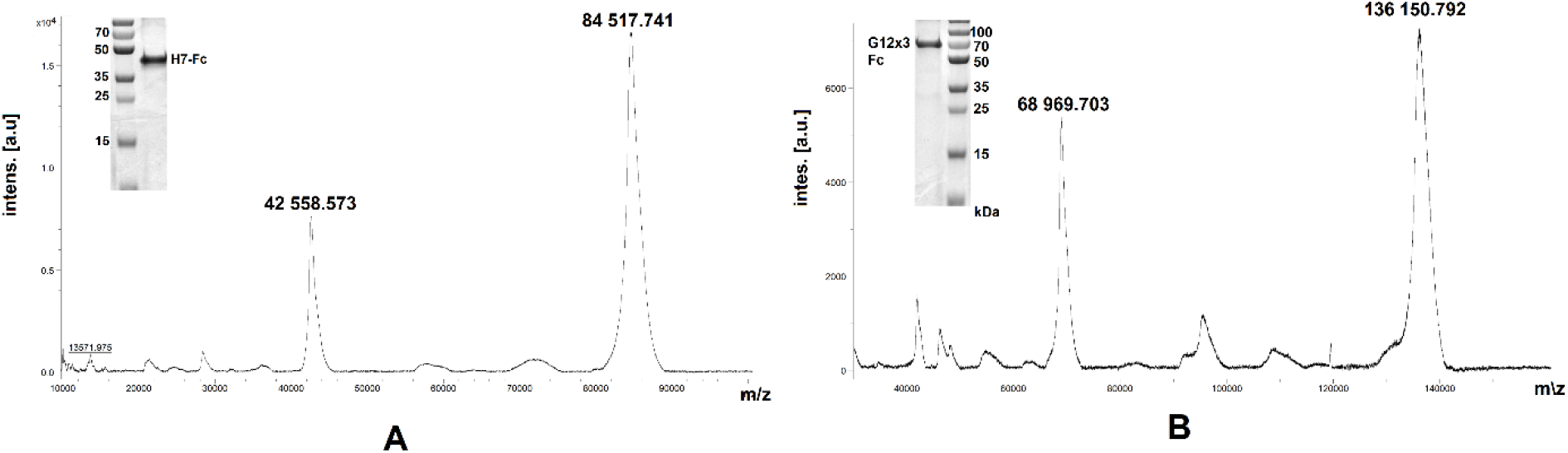
MALDI-TOF spectrum of purified G12×3-Fc (A) and H7-Fc (B) proteins. Insert: 12% SDS-PAGE reducing electrophoreses.

**Figure S3.**
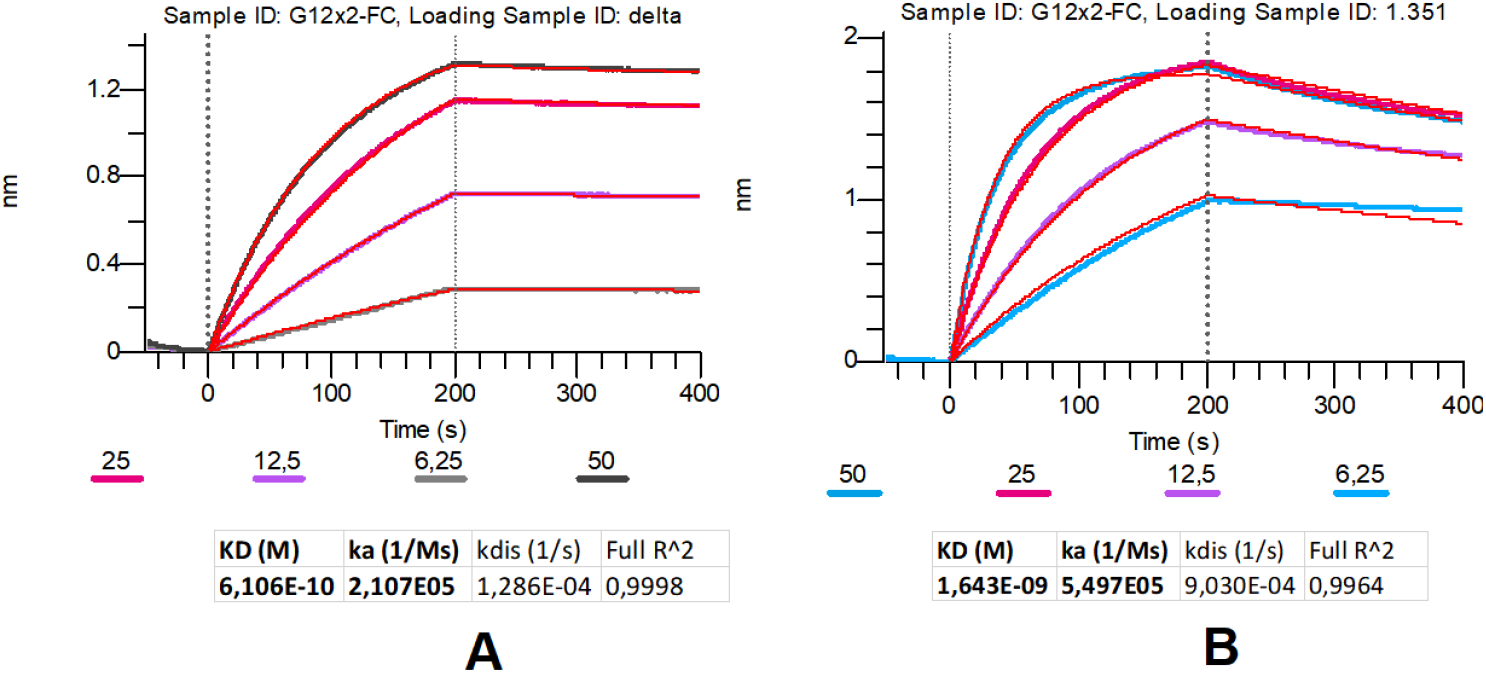
Multipoint BLI of G12×2-Fc binder to SARS-CoV-2 Delta (A) and Beta (B) RBDs.

**Table S1.** Neutralizing activity (PRNT) of top antibody candidates in comparison with the ACE2-Fc protein.

|  | IC50, ng/ml, SARS-CoV-2 variants |  |  |
| --- | --- | --- | --- |
|  | **D614G** | **Delta** | **Omicron** |
| **H7-Fc** | 133.8 | 12.3 | 106.0 |
| **G12x3-Fc** | 13.1 | 0.9 | 9.6 |
| **ACE2-Fc** | 1664.1 | 52.1 | 391.0 |

**Table S2.**
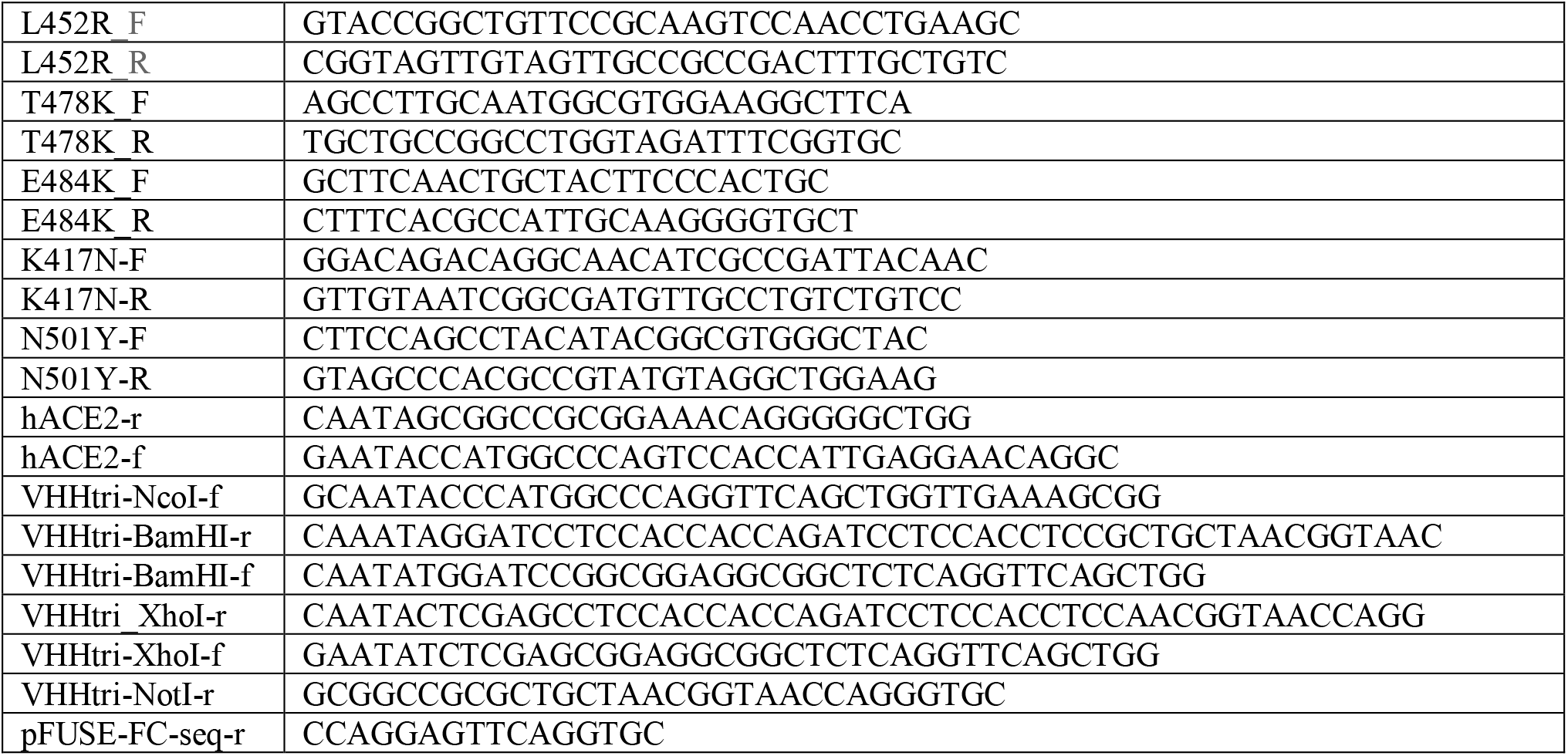
List of oligonucleotides.

